# Caring for parents: an evolutionary rationale

**DOI:** 10.1101/239285

**Authors:** J. Garay, S. Számadó, Z. Varga, E. Szathmáry

**Author notes:** Correspondence to: Eörs Szathmáry,.

## Abstract

The evolutionary roots of human moral behavior are a key precondition to understand human nature. Here we investigate whether a biological version of Fifth Commandment (“Honor your father and your mother, that your days may be long”), respected in different variants across cultures, can spread through Darwinian competition. We show by a novel demographic model that a corresponding Fifth Rule (“During your reproductive period, give away from your resources to your post-fertile parents”) will spread even if the cost of support to post-fertile grandmothers considerably decreases the demographic parameters of fertile parents but radically increases the survival rate of grandchildren. Teaching vital cultural content is likely to have been critical for the value of grandparental service. Selection on such behavior may have produced an innate moral tendency to honor parents even in situations, such as experienced today, when the quantitative conditions would not necessarily favor the maintenance of this trait.

## 1. Background

Darwin [1] already raised the possibility that moral has evolutionary origin. There are several models rooted in evolutionary theory that shed light on some basic moral issue [2–5]. In contrast, we start with a moral commandment, and investigate whether a phenotype corresponding to this moral commandment wins in a Darwinian struggle for existence or not, similar to an investigation of the conditions under which spiteful behavior will die out [6]. Here we investigate the cultural norm that promotes the help of the parents (we discuss the issue of grandfathers later). We refer to this norm as the Fifth commandment (see Supplementary Information, see SI). This norm has obvious links to biology, and variants of it can also be found in various cultures ranging from the East to the West (SI). There is widespread evidence not just for the existence of such norm but for the actual support as well. The form of this support can vary across cultures (emotional, instrumental, financial, etc.) and might be a function of other factors, such as the health of the elderly parents, but this kind of help is readily observed across different cultures [7–11]. Investigating the dynamics of such norm can shed light on the evolutionary roots of religion also [12].

In the course of standard human life history infants grow to become parents who age into being grandparents. Thus, longevities permitting, respect and help to parents turn out to be targeted to the grandparents of one’s children. This truism has important consequences for the possible spread of such a behavioral trait. Behaviors can be inherited, which can be the result of either genetic or cultural transmission. This inheritance assumption immediately implies that if the support to grandparents spreads by Darwinian selection, then that ensures longer life for the parents as their children inherit their behavior. Similarly to classical evolutionary game theory, we will not consider the genetic background of the behavior [13]. An adaptive phenotype will outperform its rivals on a Darwinian selection time scale, where Darwinian fitness is the average growth rate of a phenotype.

The establishment of a post-fertile period is critical for our case. Several hypotheses deal with the origin of the menopause.

Shanley and Kirkwood [14] investigate two alternative theories that might explain the origin of menopause. The first one can be called as the “altriciality” hypothesis that observes that maternal mortality is increasing with age. It implies a trade-off between rearing existing, still altricial children and giving birth to a new one. The second one is the mother hypothesis. The mother hypothesis states that the post-fertile grandmother helps her fertile daughter [15]. They found that neither of these ideas alone is sufficient to explain the evolution of menopause under a realistic range of life-history parameters; however, a combined model can explain it [14, 16]. Their conclusion is corroborated by other studies both with regard to altriciality [17] as well as to kin-selection [18].

According to the grandmother hypothesis [19–24] the advantage of the post-fertile stage is that grandmothers increase the survival of their grandchildren [20, 24, 25]. increasing either the survival rate or the fecundity of the latter [26, 27]. A third hypothesis is the “embodied capital model”, which emphasizes that the inter-generational transfer of skill, knowledge and social ability needs time, and both grandmothers and grandfathers could help the “training” of their grandchildren [28]. The attained skills and knowledge during childhood can increase the survival rate and fecundity for the whole adult life period of the grandchildren; see Figure 1.a-c for a comparison of these alternatives. These three hypotheses do not necessary exclude each other since the care for pre-fertile individuals includes breastfeeding, transport, feeding and protection as well as affection and education [25, 29, 30].

**Figure 1.**
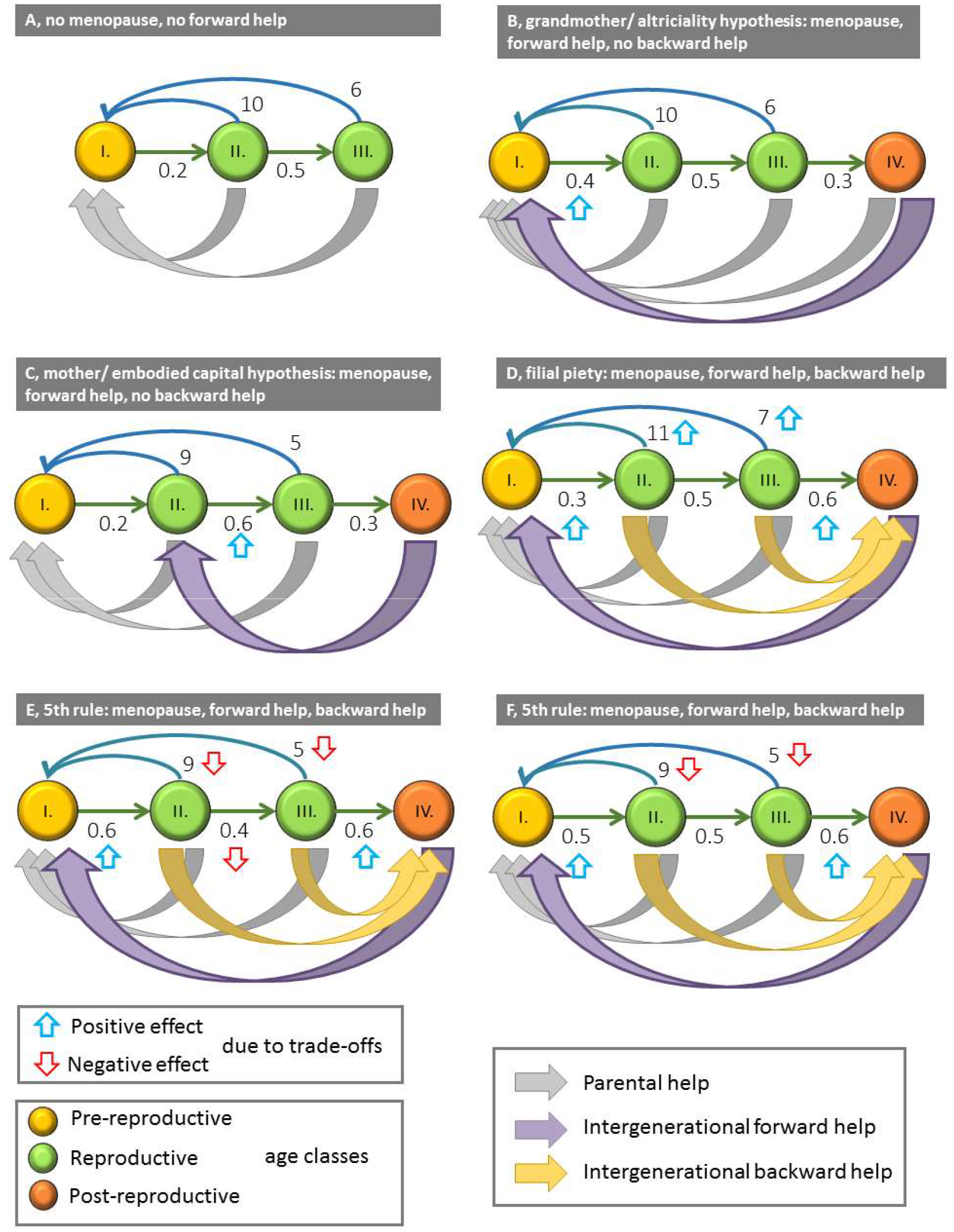
Schematic representation of the different theories. Grey arrows denote parental help; purple arrows denote forward help from the grandmothers to the grandchild, finally yellow arrows denote backward transfer of resources from the parents to the grandparents. Upward blue and downward red arrows denote positive and negative effects from trade-offs respectively. (a) Standard life-history model, no menopause, no forward help and no backward help; (b) grandmother (purple arrow from VI to I)/ altriciality (grey arrow from VI to I) hypothesis: menopause evolved, no backward help from parents to grandmothers; (c) mother and/or embodied capital hypothesis: menopause evolved, no backward help from parents to grandmothers; (d) filial piety: menopause evolved, synergistic division of labor with backward help from parents to grandmothers, no trade-offs; (e) Fifth rule: menopause evolved, backward help from parents to grandmothers, three-way trade-off for the parents between survival, fecundity and helping the grandmothers; (f) Fifth Rule: menopause evolved, backward help from parents to grandmothers, two-way trade-off for the parents between fecundity and helping the grandmothers.

All these hypotheses are aimed at explaining the evolutionary advantage of the long post-fertile life period of Homo *sapiens.* However, none of them assumes a transfer of resources from the parents to the grandparents, thus none of them investigates the trade-off situation between parental reproduction or survival and the support to grandparents. The central question is this: Will support to post-fertile grandmothers spread even if there is a trade-off between this support and either the fecundity or the survival rate of fertile parents?

Chu & Lee [3] investigated the evolution of intergenerational transfer (IT) from parents to grandparents in the framework of a cooperative game and they already pointed out that “filial piety” can evolve by division of labor. Fertile female transfers some energy to her mother, enabling the latter to redirect her efforts from inefficient foraging to grandchildren care. During this time the fertile female is free from caring and she can go to forage with higher efficiency than her mother. In other words, this model describes a synergistic situation where everyone does the task she is the most efficient in. But the authors do not consider the trade-off we wish to investigate (see Figure 1.d).

We strongly concur with the statement that “Even to demonstrate, for example, that post reproductive women result in a reduction in grandchild mortality *does not establish that menopause is adaptive unless it can be demonstrated that overall fitness is actually enhanced”* [16] (pp. 27, their emphasis). In establishing the selective advantage of care for grandmothers we consider the effect of overall fitness of the family.

Since in our problem pre-fertile, fertile and post-fertile individuals live together in a family, we have to consider a kin demographic selection model [3, 31, 32], in which the survival and the fecundity parameters depend on the costs and benefits of intra-familiar supports. After setting up the model we investigate whether the Fifth Rule (as a biological distillation of the Fifth Commandment; see *Materials and Methods)* wins in a Darwinian struggle for existence. Finally, we discuss our results.

## 2. Results

We consider a Leslie matrix model (see *Materials and Methods).* What is the effect of the Fifth Rule on the entries of the above Leslie matrix? For the simplest mathematical formulation we assume that the cost of supporting grandmothers does not depend on the age class of either parents or grandmothers. Let *y* ∈ [0,1] be the cost spent on grandparent support. If grandmothers help in child care, the survival rates of children *ω_i_* increase withy, and based on the grandmother and the mother hypotheses, *ω_j_* decreases and *α_j_* increases with increasingy, where *ω_i_*(*y*) (*i* = 1,…,*k*) denote the survival rates of children, *ω_j_*(*y*) and *α_j_*.(*y*) (*j* = *k* + 1,…,*K*) are the survival rate and fecundity of fertile parents, respectively.

Since there is a difference in intra-familiar support between families, the Leslie matrices of different family types are different. What kind of intra-familiar support ensures the highest long-term growth rate for the family? For sake of simplicity we denote help from the grandmothers to children as “forward help” and help from the parents to the grandparents as “backward help”. (See Fig.1 for comparison of the different models.) Under well-known conditions (fulfilled in our case), the unique positive eigenvalue of Leslie-matrix is the longterm growth rate of the family, thus we consider this eigenvalue as fitness [33, 34]. Formally, the fitness *λ*(*y*) is the unique positive eigenvalue of the y-dependent Leslie matrix (see *Materials and Methods*), hence, other things being equal, families helping grandmothers are competitively superior to those without this behavior.

### Grandmother hypothesis

Consider the case when fertile individuals do not support grandmothers (see Figure 1.e for a general depiction of the idea). We consider the following two cases (see *Matherials and Methods):* (i) If grandmothers do not help in child care, but their survival linearly reduces their own fecundity, then the optimal strategy is not to spend on own survival to post-fertile age. (ii) If grandmothers help in child care then the menopause is evolutionarily successful if the effect of grandchild care 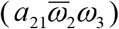 on the grandchild’s survival 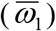 is greater than his/her survival rate without this care, i.e. 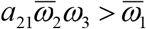 (see Table 1 for notation, 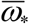 denotes averages). Summing up, the grandmother hypothesis concerns the way a female of reproductive age allocates her resource between own survival and fecundity. Note that we have adopted the hypothesis that the cost spent on living to the post-fertile age reduces fecundity. Without this trade-off, living to the post-fertile age is a neutral property in the first case, and a benefit in the second case.

**Table 1.**
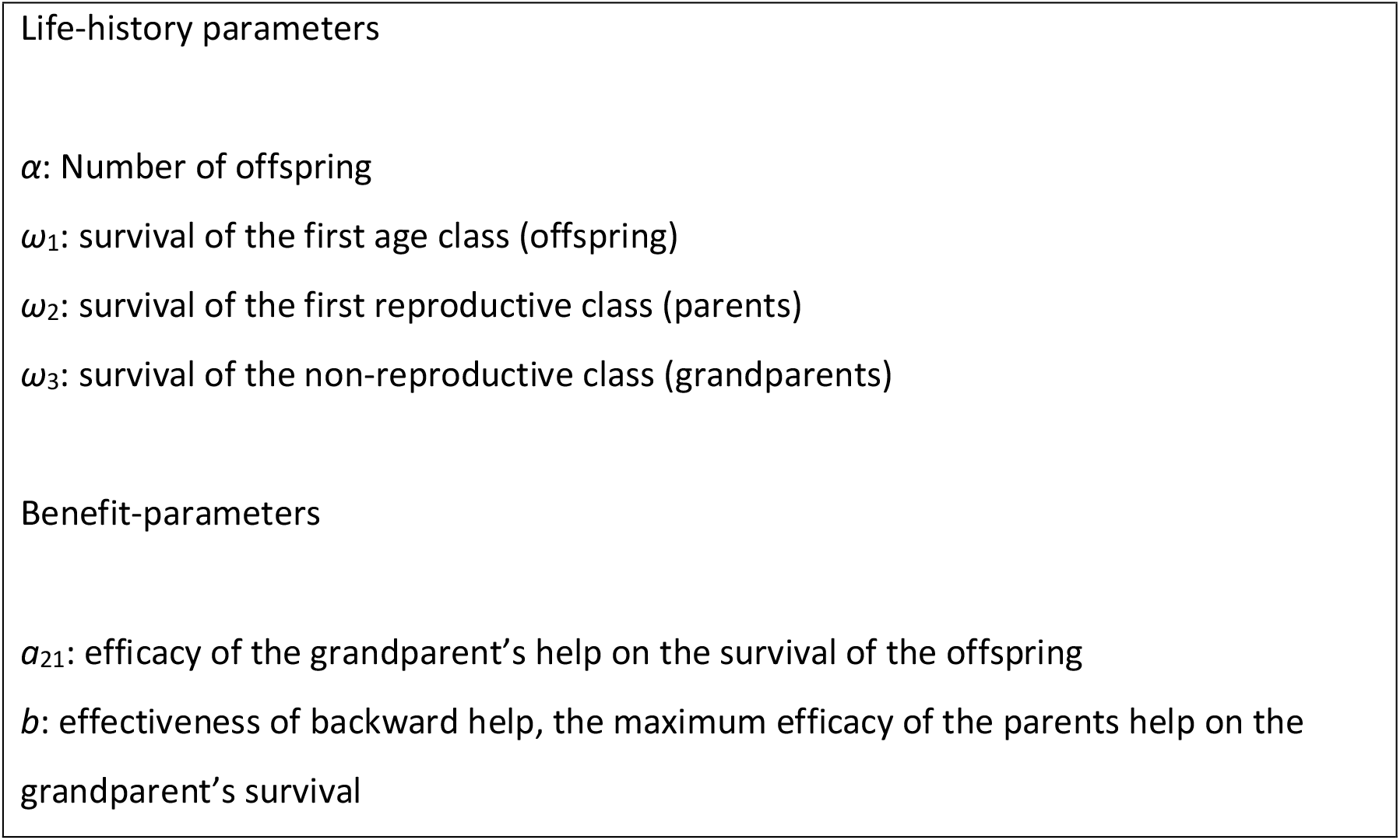
Notation of the model.

### The Fifth Rule

The fifth rule requires us to support our elderly (see Figure 1.f for a general depiction of the idea), which may occur when the menopause has already become evolutionarily fixed (see *Matherials and Methods).* We show (SI) that the Fifth Rule (backward help) evolves when 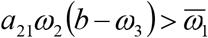. This condition is satisfied if, for example, the efficiency of the support to post-fertile parents is sufficiently large compared to the basic post-fertile survival rate (if the latter is high than grandmothers would be around even if not helped).

Of course, the coevolution of two traits: long life in menopause and an effective Fifth Rule is also possible. The analytical study presented in *Materials and Methods* is based on two conditions. First, the development of the Fifth Rule is conditional on the existence of the menopause, since one can help a grandmother only if she is alive. (Based on this conditionality, in the *Materials and Methods* we suppose that traits *s* and *y* determine both the increase of the grandmother’s survival probability and the decrease of fecundity in multiplicative form.) Consequently, the rarity of an effective Fifth rule is hardly surprising. Second, if a fertile mother were to give away all her resources to help the survival of her mother, her fecundity drops to zero. In the *Materials and Methods,* in terms of a fitness landscape, we show that, if the fitness *λ*(*s, y*) has a global strict maximum point (*s*,y\**) (e.g. in our case if *λ*(*s, y*)) is strictly concave), then there exists an unique evolutionarily optimal behavior (*s*, y\**), hence the species evolves into this state.

The Fifth Rule will spread if the cost of support to post-fertile grandmothers decreases slightly the demographic parameters of fertile parents, but sufficiently increases the survival rate of grandchildren. However, in general, there is a threshold over which support to grandmothers has no evolutionary advantage. If the cost of support to post-fertile grandmothers only decreases the demographic parameters of the family, but it offers no increase in the survival rate of the grandchildren, then the Fifth Rule has no evolutionary advantage. The mother hypothesis and “embodied capital model” should imply that grandmothers increase the survival rate of their children and that of grandchildren during their lives. Thus, if these ideas also work in human evolution then it is even “easier” for the Fifth Rule to evolve (see *Materials and Methods*).

In order to investigate the effects of different cost-benefit parameters on the evolvability of IT we have constructed a general example, which was analyzed numerically (see *Matherials and Methods).* Conclusions of the model are as follows: IT evolves most readily when the grandparental help increases both the survival and the number of offspring [20, 24, 25] (Figure 2, Figs. S1-S3). Linear cost and benefit functions do not favor the evolution of IT (Figs. S1, S4, S6); conversely, convex benefit and concave cost functions promote the evolution of IT (Figs. S2, S3, S5, S7). It is possible to find cost parameters (*c,d*) where IT evolves even if the efficacy of parental transfer and grandparental help (*a*_21_ and *b* respectively) is low (Figs. S2, S3). Conversely, it is possible to find (high) a21, *b* parameters where IT evolves even if it imposes a high cost on the survival of the parents or on the number of offspring (*d* and *c*, respectively, see Figs. S1, S2).

**Figure 2.**
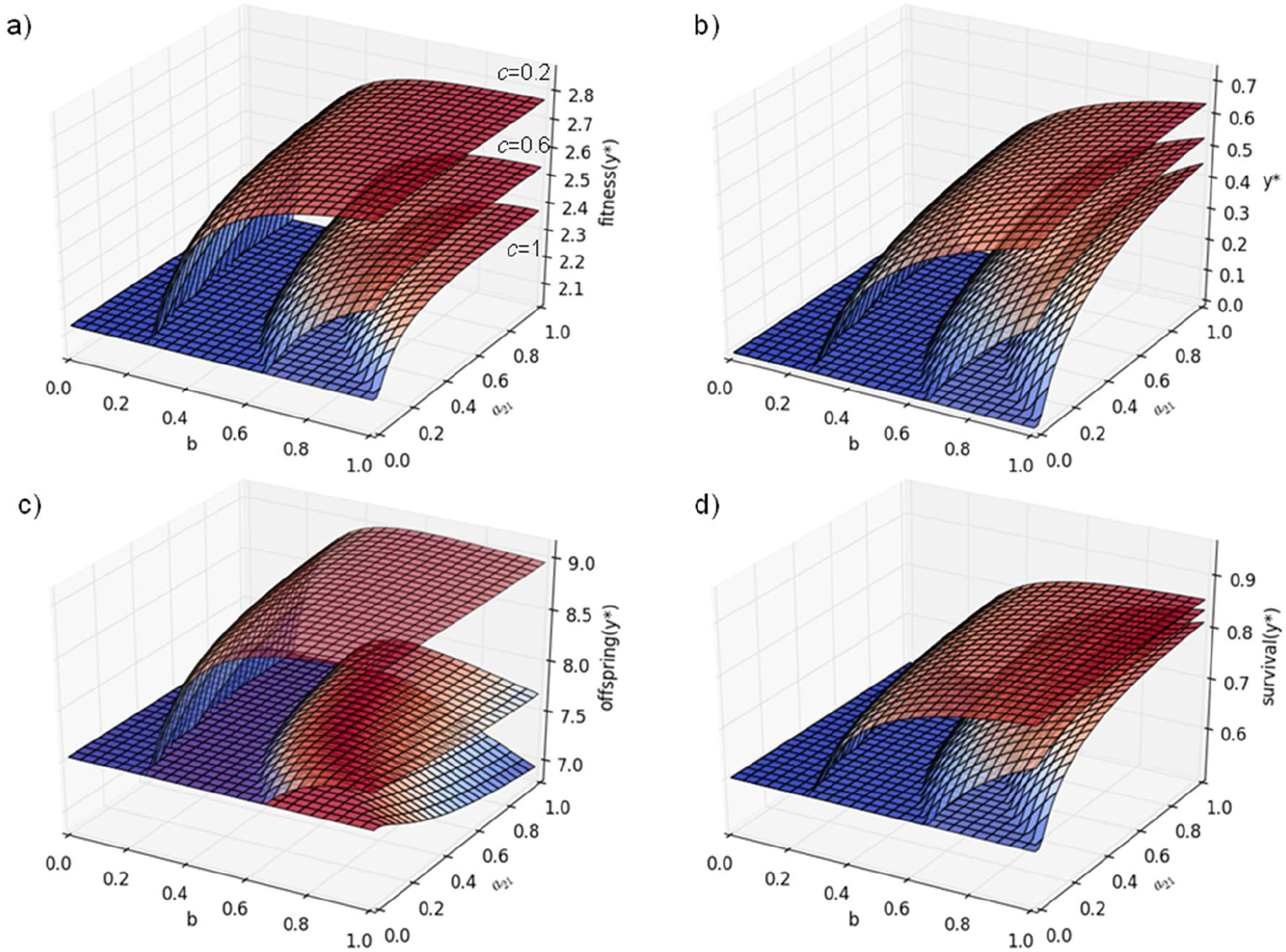
Numerical example for a 2x2 Leslie matrix (see *Materials and Methods* for details). (a) Maximum family long term growth rate (fitness); (b) optimal level of backward help (*y*\*), (c) average number of offspring at *y*\*; (d) offspring survival at *y*\* as a function *b* (effectiveness of backward help) and *a*_21_ (effectiveness of forward help to offspring survival). Parameters: *α*_2_=6, *ω*_1_=0.45, *ω*_2_=0.62, *ω*_3_=0.25, *d* = 0.3, *h* = 1, *c* =0.2, 0.6, 1; *a*_12_=10.

Since we are dealing with “family issues” the natural conceptual framework is that of kin selection. Our problem does not, however, readily yield to standard inclusive fitness modeling, since the latter is not sensitive to demography. There are different contributions to a female’s fitness from the three stages of her life-history: girl, mother and grandmother. Our demographic model can account for these complications in a straightforward manner. Note also that our account involves an unusual loop from parent to grandparent to grandchild. Our analysis applies to grandfathers as well, provided the menopause in grandmothers constrains the realized fertility of the former in a similar way.

## 3. Conclusions

We demonstrate that an essential part of Fifth Commandment (support to elderly) can confer selective advantage under the right conditions; hence some kind of “evolutionary moral sense” might be genetically endowed. This holds if grandparents have a positive effect on the growth rate of their family. However, this is not necessarily true nowadays [35]. It is very well possible that this support can be rooted in past human evolution. Darwinian success of the Fifth Rule cannot completely explain the present-day Fifth Commandment. Human moral rules, although rooted in Darwinian evolution, are more than what that theory supports. It seems that the main difference is that moral commandments are unconditional rules, while in Darwinian evolution there must be a selective condition determining whether a behavior is adaptive or not.

## 4. Materials and Methods

### 4.1 Cultural analogues of the Fifth Commandment

Cultural norms that promote the help of the parents are widespread in both western and eastern culture. The Fifth Commandment (of the Hebrew and protestant Bible, the Fourth one, according to the catholic numbering) states: *“Honor your father and your mother, that your days may be long in the land that the LORD your God is giving you*.” (Exodus 20:12) From the interpretations of this commandment by the western churches we recall the following: Sefer Ha-chinukh (mitzva 33) elaborates: “*A person should realize that his father and mother are the cause of his existence in this world; therefore it is appropriate that he render them all the honor and do them all the service he can*” St. Thomas Aquinas wrote: “*Since we receive nourishment from our parents in our childhood, we must support them in their old age*.” Martin Luther said: “*For he who knows how to regard them in his heart will not allow them to suffer want or hunger, but will place them above him and at his side, and will share with them whatever he has and possesses*” (Luther, M. p. 29).

We also note that in China, to take care of elderly parents is also a moral rule: e.g. Confucius declared: “*In serving his parents, a filial son reveres them in daily life; he makes them happy while he nourishes them; he takes anxious care of them in sickness* …” (26) Based on the above, we introduce the so-called Fifth Rule, which is a translation of the Fifth Commandment into biological terms and is inherent in the above interpretations: “During your reproductive period, give away from your resources to your post-fertile parents.”

### 4.2 The model

Our model strictly follows the Darwinian view: the fitness is determined by the fecundity and the survival rate. The fecundity of the family is determined by the intergenerational help, which modifies the demographic parameters within the family. Furthermore, the carrying capacity also has an effect on the survival. Thus, the survival of an individual depends on the intra-familiar help, see subsection 4.3, and the survival probability according to the carrying capacity, see subsection 4.4. Our model combines these two factors.

### 4.3 The phenotype-dependent Leslie matrix

We consider the following age-structured model with two sub-models. The development of a family is described by the following Leslie matrix, which contains the survival and fecundity parameters of pre-fertile and fertile individuals, and all entries depend on the level of the intra-familiar (backward) help, denoted by *y*:

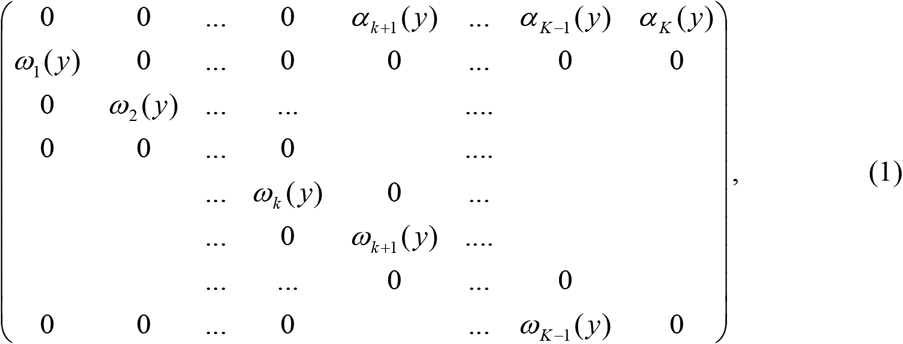

where *ω_i_*(*y*) (*i*=1,…,*k*) denote the survival rates of children, *ω_j_*(*y*) and *a_j_*(*y*) (*j*=*k*+1,…,*K*) are the survival rate and fecundity of fertile parents, respectively. Figure SI1 depicts an example. The multiplication of the age-structured population vector by this Leslie matrix describes the dynamics of the family. The age classes of grandparents will be handled separately, since the development of the family depends on the survival rate of pre-fertile and the fecundity of the fertile family members. Formally, *x_l_* = *ω_l-1_* (*y*)*x_l-1_* where *ω_l_* (*y*) (*l*=*K*+1,…,*H*) are the survival rates of the grandparents, and *x_l_* is the number of grandparents in age class *l.*

### 4.4 The survival at the carrying capacity

In the framework of the Leslie model, it is widely accepted that the Darwinian fitness is the long-term growth rate of the phenotype (i.e. the dominant positive eigenvalue of the Leslie matrix, see *23*), surprisingly, in the literature we could not find a Darwinian explanation to this. Below, adapting our recent reasoning from (27), we propose a strictly Darwinian reasoning to see that the long-term growth rate is maximized by natural selection: The number of offspring, in general, is much higher than the carrying capacity, so only a part of the offspring and adults will survive. Let us consider random survival, assuming that the survival probabilities of individuals do not depend on phenotypes (in our case intergenerational help) and on the age of individuals. (Observe that this assumption gives some advantage to the families in which the intergenerational help is less.)

Now let us consider two phenotypes A and B with respective long-term growth rates (i.e. positive eigenvalues of the corresponding Leslie matrices) *λ*_A_, *λ*_B_ > 1 with λ_A_>λ_B_. To see the asymptotic frequency of phenotype B, we suppose that phenotypes A and B start from respective initial densities *x*(0) and *z*(0). According to the original Darwinian view, we need some density dependent selection to keep the total density of these two phenotypes at the carrying capacity. Since in the considered selection situation there is no interaction between the phenotypes and we assume that the phenotypes differ only in the demographic parameters, thus we can suppose there is a uniform survival process, i.e. the survival rate corresponding to the carrying capacity is the same for all individuals. Now the question arises which phenotype will win in the struggle for existence on the long selection time scale?

Let us suppose that phenotypes A and B develop according to Leslie models having the respective population vectors *x*(*t*), *z*(*t*), and matrices *L_A_*, *L_B_*, total densities 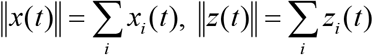. Then the relative frequency of phenotype B tends to zero, as it is shown below:

Indeed, let us suppose that the subpopulations start from initial states *x*(0) and *z*(0), respectively, and the time unit is chosen in such a way that in unit time the total density of the system always exceeds the carrying capacity *K*, in particular

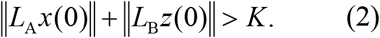

Now, by the selection the total density of the system is reduced to *K* proportionally:

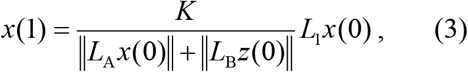

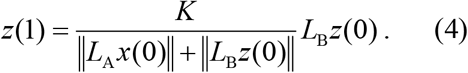

Indeed, obviously ||*x* (1)|| + ||*z*(1)|| = *K*.

We emphasize that in this model we consider the “intrinsic” survival (described by the Leslie matrices) and the survival under selection independently. However, this model can be formally considered as a particular Leslie-type model depending on the total density of the system, where each demographic parameter in the Leslie matrices *L_A_* and *L_B_* is multiplied by 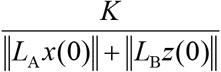.

Similarly, for all *t*= 1, 2, 3,… we get our kin demographic selection model for two different phenotypes:

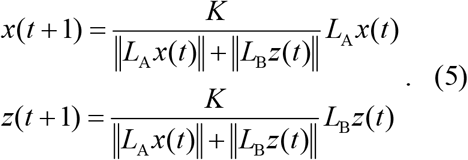

Now, for the proportion of phenotype B we obtain

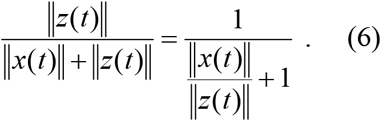

Here

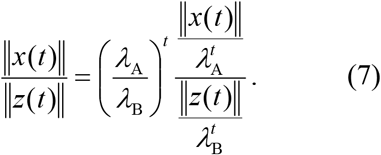

Since we can suppose that in both phenotypes the last two fecundities are positive, so the Perron-Frobenius theorem (see e.g. 28) implies that both 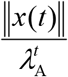 and 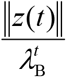 tend to finite positive limits as *t* → ∞. In fact, the Leslie matrices can be cut at the last fertile age class, apply the Perron-Frobenius theorem to these matrices, and then the convergence can be extended to the post fertile age groups by simple survival Therefore, 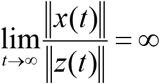 implying

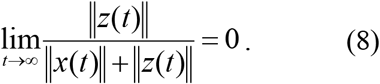

Thus if *λ*_A_ > *λ*_B_, then the relative frequency of phenotype B tends to zero as *t* tends to infinity. Observe that in our model, the fecundity of a phenotype is determined by a phenotype-dependent Leslie matrix, and the survival rates corresponding to the carrying capacity of different phenotypes are the same, so the long-term growth rate of a phenotype determines the fitness.

### 4.5 The general results

Consider the general *K* x *K* Leslie matrix, where the entries depend on the cost *y* spent to grandparent support. Under the grandmother hypothesis, the grandmother support decreases the fecundity and survival rate of fertile parents, but increases the survival rate of the grandmother, who therefore increases the survival rate of pre-fertile grandchildren. Then the characteristic equation is

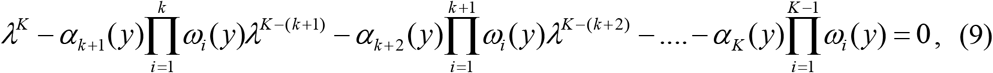

and its unique positive root is obtained as the root of equation

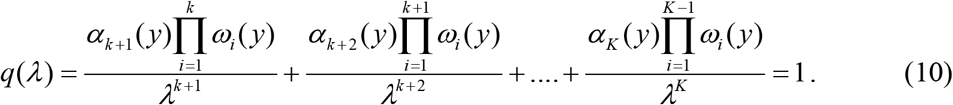

It is easy to see, that if any of the numerators (i.e. the average numbers of offspring produced by an individual of the corresponding age classes) in these fractions is changed to a greater one, then the curve of the ‘hyperbolic’ function *q* shifts upwards, implying that the positive solution *λ*_*_ of this equation also will be greater. Therefore, if in a population where within the families grandparents are not supported, a new type emerges which supports grandparents, and all mentioned numerators increase, then Fifth Rule as behaviour type will propagate. If all these numerators decrease then this type will die out. Those mathematical cases when some of the numerators increase, others decrease, would need further mathematical discussions.

Observe that equation *q*(*λ*) = 1 can be written as

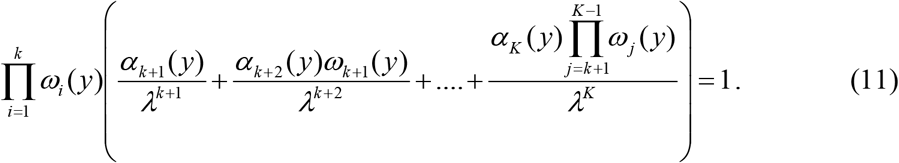

Here factor 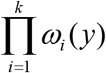 measures how much child care by grandmothers increases the survival of the children. Roughly speaking, factor

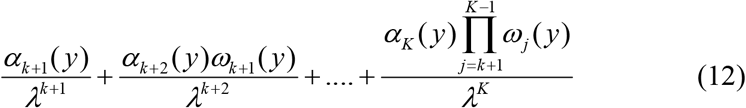

measures, in an implicit way, to what extent the support to grandparents by the fertile age class decreases their own fecundity and survival rates. In this sense, the predictions of our model are in harmony with the cost-benefit approach saying that a trait will propagate if it eventually increases the fitness.

Now the question arises how the demographic parameters may depend on *y*. The following assumptions are at hand: 1. The survival rate of grandparents is a saturation function of *y* strictly increasing at the beginning, and remains constant after. 2. Based on the grandmother hypothesis, the survival rate of grandchildren strictly increases with the survival rate of grandparents (which on its term depend on *y*). The grandmother hypothesis is the worst case when two trade-offs may exist. 3. The parents’ fecundity entries of the Leslie matrix (*α*_*k*+_,…,*α_K_*) are strictly decreasing functions of *y*. 4. The survival rates of parents (*ω*_*k*+1_,…,*ω_K_*) are strictly decreasing functions of *y*.

These assumptions allow the Fifth Rule to win or lose the struggle for existence, depending on whether the long-term growth rate of the family increases or decreases.

Under assumption 1 there is a threshold for the support to grandparents, above which the survival of grandparents does not increase, and therefore the survival of grandchildren either, but the fecundity and/or the survival of fertile parents still decrease. Over this threshold, the support to grandparents has no evolutionary advantage.

Finally, we remark that the above reasoning can be applied not only to the grandmother hypothesis, since either the mother hypothesis or the embodied capital model alone can ensure the support to grandparents. For example, if any of the above two hypotheses implies the increase of at least one of the numerators in (3), while the rest of the numerators do not decrease, then the dominant eigenvalue, i.e. the asymptotic growth rate will increase. Of course, if in addition to the fact that the grandmother increases the survival of her grandchildren and the survival and fertility of her daughter, the hypothesis of the embodied capital model also holds (the grandmother also increases the adult age survival and fertility of her grandchildren), then not only 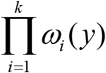 but also factor (4) can synergically increase the asymptotic growth rate of the family.

Finally, we note that the “altriciality” hypothesis can also be handled in terms of a linear model with a matrix structured differently from the Leslie matrices (since the survivals of children also depend on the age of their mothers). Thus, only a generalization our model could deal with the development of menopause based on altriciality. In our opinion, our Fifth Rule may be derived on the bases of “altriciality” hypothesis, but in such a future model the formation of multi-generation families should also be included, since “altriciality” hypothesis itself does not need the convivence of several generations.

### 4.6 Two-age-class model

For a deeper insight, in this simplest case, we will calculate first when the menopause can evolve, second, when the Fifth Rule is evolutionary successful, third, using numerical examples we demonstrate that convex benefit and concave cost functions promote the evolution of intra-familiar help.

Consider the simplest case with one child age class and one fertile age class. Then the Leslie matrix is

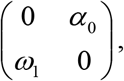

where *ω*_1_ is the survival rate of children and *α*_0_ is the fecundity of fertile parents. The survival rate from fertile age to post-fertile age is *ω*_2_, and *ω*_3_ denotes the probability that a post-fertile individual still lives (without the support by a fertile individual) when child care is needed. Now the fitness is 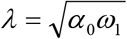.

### 4.7 Grandmother hypothesis

Now the question arises: When is the menopause adaptive? Consider the case when fertile individuals do not support grandmothers. We consider the following two cases: (i) Suppose that grandmothers do not help in child care, but their survival linearly reduces their own fecundity, i.e. *ω*_2_(*s*) ≔ *s* and *α*(*s*) ≔ *α*_0_(1 –*s*), where *s* ∈ [0,1) is the cost spent on survival to post-fertile age (Figure 3.a depicts the situation). The fitness of the population is the long-term growth rate which can be calculated from the characteristic equation of the Leslie matrix: 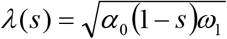, and the optimal strategy is not to spend on own survival to post-fertile age. (ii) Suppose that grandmothers help in child care (Figure 3.b). Let 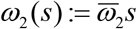, with some 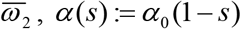 and *ω*_1_ (*s*):= *ω*_1_ + *a*_21_*P*(*s*), where *s* ∈ [0,1), *ω*_1_ is a "basic” survival rate, and the probability that a grandmother is alive when her help needed is 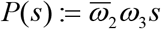, i.e. we count only the help of those grandmothers who survive to the upper boundary of the third age class and do not count those who reach ‘grandmother age’ (reach the third class) but die before the upper boundary of age, and *a*_21_ denotes the efficiency of the grandmother’s grandchild care. Clearly 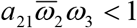 and 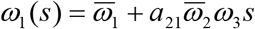, thus the fitness is

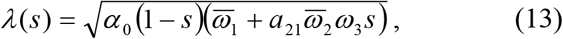

which is maximal at 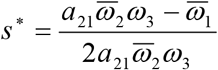. Therefore, if the effect of grandchild care on the grandchild’s survival is greater than his/her survival rate without this care, i.e. 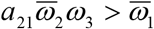, then menopause is evolutionarily successful.

**Figure 3.**
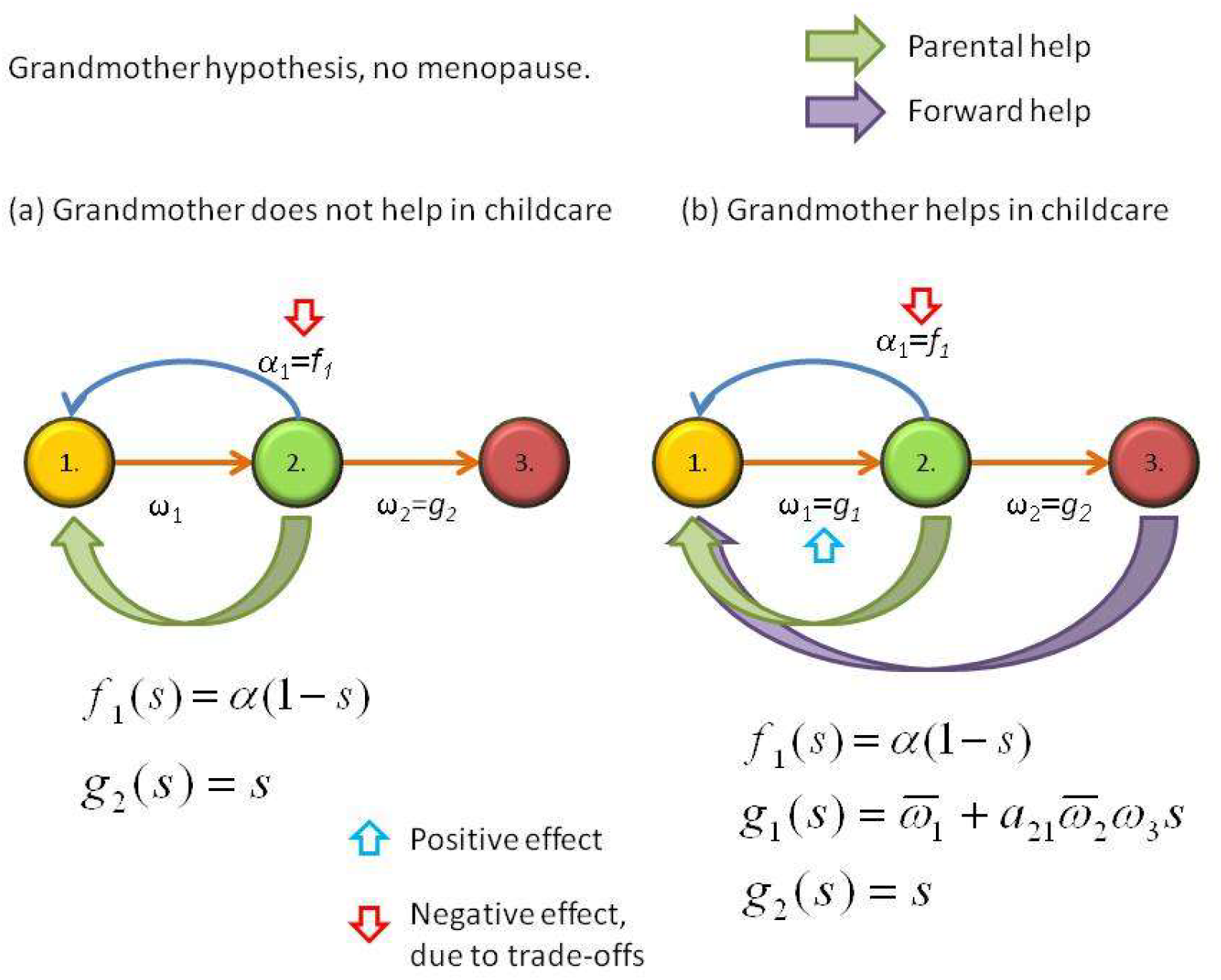
Grandmother hypothesis without (a) and with (b) childcare. Green arrows denote parental help; purple arrows denote forward help from the grandparents to the grandchild.

### 4.8 The Fifth Rule

Now the question arises: When is Fifth Rule adaptive? It requires us to support our elderly, which is possible only if the menopause has already become evolutionarily fixed, i.e. for fixed *s* ∈ [0,1), let 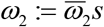 and *α*: = *α*_0_(1 – *s*). Let *y* ∈[0,1] denote the cost spent on the survival of post-fertile parents, and suppose that the negative effect of *y* on fecundity is linear: *α*(1 – *y*), the children survival is 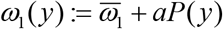, where *P*(*y*) ≔ *ω*_2_(*ω*_3_ + *by*) and *b* indicates how efficiently the support to post-fertile parents by fertile individuals increases post-fertile survival, so 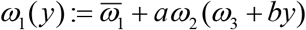 (Figure 4 depicts the situation). Now we have to maximize the fitness which can be calculated from the characteristic equation of the Leslie matrix, it is given by the following function in *y*:

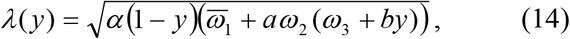

which attains its maximum at 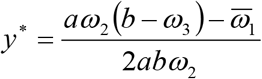. The latter is positive if 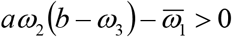. This condition is satisfied e.g., if the efficiency of the support to post-fertile parents is sufficiently large compared to the basic post-fertile survival rate.

**Figure 4.**
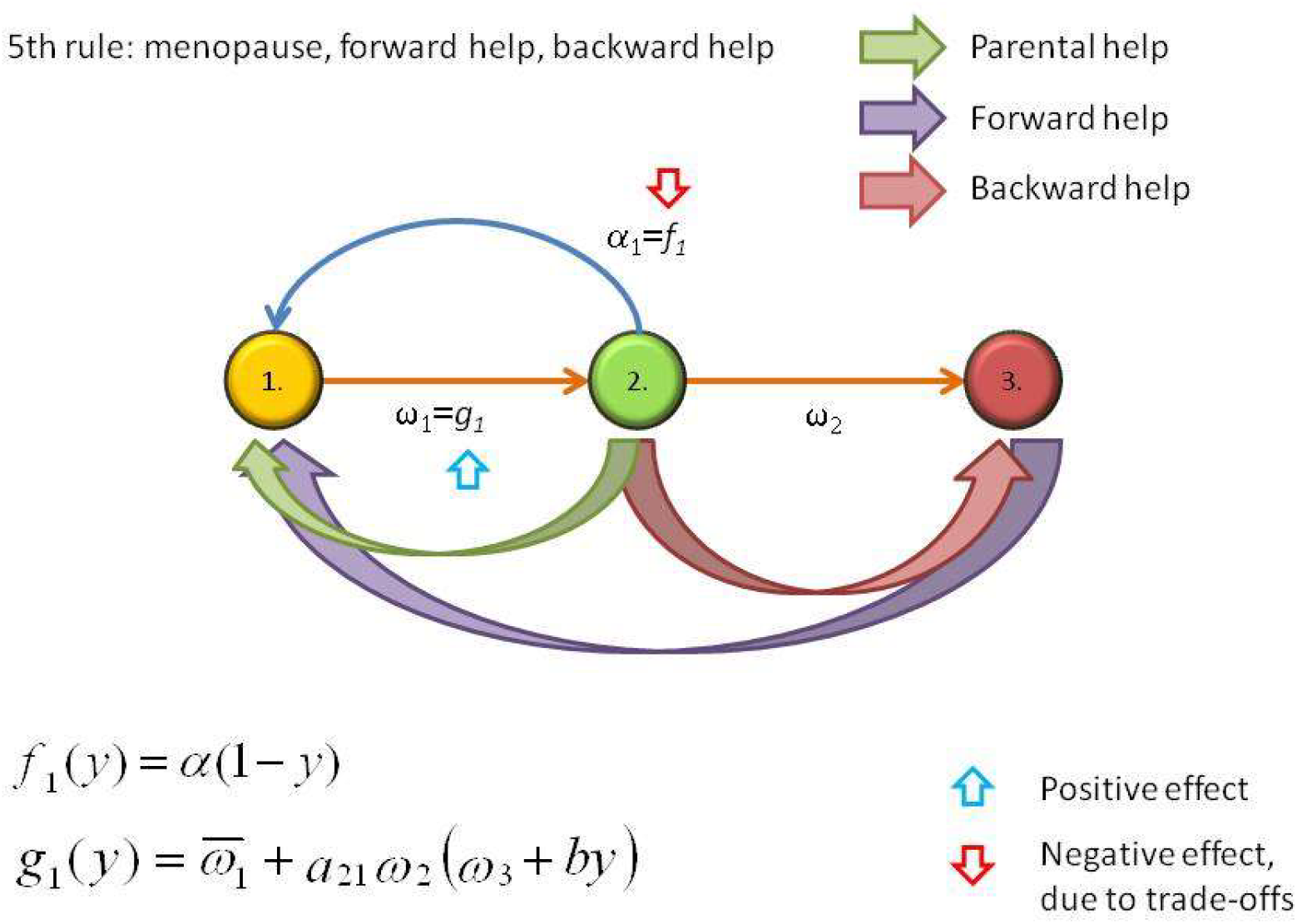
The fifth rule: forward help in childcare and backward help to the grandparents. Green arrows denote parental help; purple arrows denote forward help from the grandparents to the grandchild, finally red arrows denote backward transfer of resources from the parents to the grandparents.

### 4.9 A general multiplicative coevolution model

Now we set up a model combining the model of grandmother hypothesis and the model of the Fifth Rule. Our study will be based on two biological preconditions: First, since one can help a grandmother only if she is alive, for the development of the Fifth Rule, the existence of menopause is needed. Second, if a fertile mother gave away all her resources to help the survival of her mother, her fecundity would be zero. As before, let *s* be the cost a fertile female spends on her own survival to post-fertile age, and *y* the cost a fertile female spends on the survival of post-fertile parents. Based on the first precondition, unlike the additive approach of sections 3.2 and 3.3, we express the effect of strategies s and *y* on the demographic parameters in multiplicative form, considering the following strategy-dependent Leslie matrix:

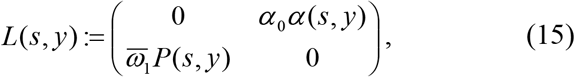

where in both variables *P*(*s, y*) is strictly monotonically increasing, and *α*(*s,y*) is strictly monotonically decreasing. Let us assume that strategies s and *y* act independently both on the fecundity and on the survival of children:

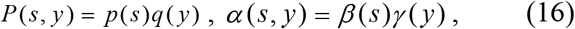

where all one-variable functions are defined on [0,1].

Technical conditions on the functions involved:

a. *p, q, β, γ* are twice continuously differentiable.
b. *p*(0) = *q*(0) = 1, *β*(0) = *γ*(0) = 1. We note that these technical conditions imply that 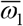 and *α*_0_ are the demographic parameters before the appearance of the considered traits, while *β*(1) = *γ*(1) = 0 expresses our second precondition.
c. *P′(s) q′(y)* > 0 (*s,y* ∈ [0,1]), 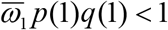.
d. *β′(s)*, *γ′(y)* < 0 (*s, y* ∈ (0,1]), *β′*(0) = *γ′*(0) = 0. Observe that conditions c) and d) are mathematical descriptions of trade-offs.
e. *p″(s)*, *q″(y),β″(s)*, *γ″(y)* < 0 (*s, y* ∈ (0,1)). (This condition will guarantee strict concavity of function z near its maximum).

Now, the fitness (unique positive eigenvalue of *L*(*s, y*)) is

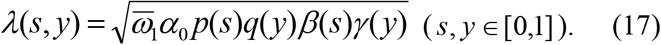

We will show that *λ*(*s, y*) attains a strict local maximum at an interior point of the unit square [0,1] x [0,1]. Indeed, maximization of *λ*(*s, y*) is equivalent to the maximization of

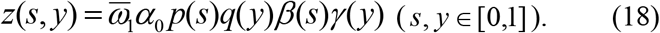

The first order necessary condition for the maximum attained at an interior point is

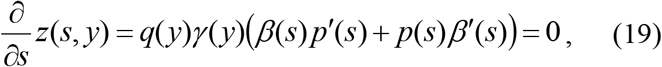

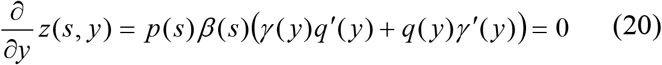

Since *p, q, β, γ* are all positive in the interval (0,1), the above necessary condition is equivalent to

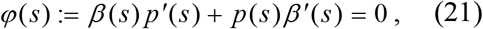

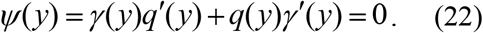

From conditions b), c) and d) we obtain *φ*(0) > 0, *φ*(1) < 0, hence there is an *s*\* ∈ (0,1) with *φ*(*s*\*) = 0. It is easy to check that conditions b), c), d) and e) also imply *φ*′( *s*) < 0, and hence *φ* is strictly decreasing, therefore *s*\* is its unique zero in the interval. (0,1). Similar straightforward checking shows that *ψ*(*y*) also has a unique zero *y*\* in the interval (0,1). Hence (*s*\*,*y*\*) is a unique stationary point of function *z* in the interior of the unit square. Now, for a second order sufficient condition for the maximum of function *z*, we calculate its Hessian:

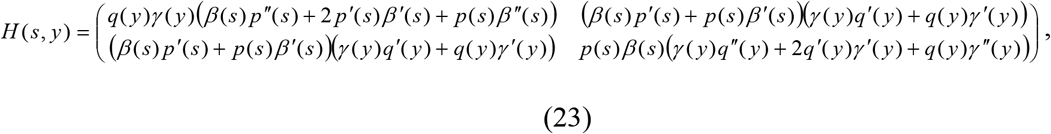

Observe that from *φ*(*s*\*) = 0 and *ψ*(*y*\*) = 0, we obtain

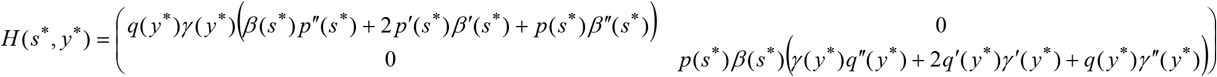

From conditions a)-e), we easily get

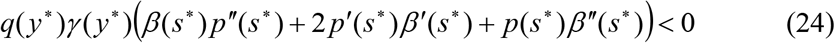

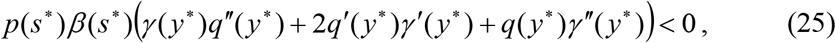

implyig tr *H* (*s*\*, *y*\*) < 0 and det *H* (*s*\*, *y*\*) > 0, i.e. *H*(*s*\*, *y*\*)is negative definite. Therefore (*s*\*,*y*\*) is a strict local maximum point. Since (*s*\*,*y*\*) is the unique stationary point, it is also a strict global maximum point in the interior of the unit square.

Finally, we note that, if Hessian *H* (*s, y*) is negative definite in the interior of the unit square, then function *λ* is globally strictly concave, and therefore (*s*\*, *y*\*) is a global maximum point of *λ*. In the terminology of fitness landscapes, in the sense of any reasonable strategy dynamics the species will evolve into the evolutionarily optimal behavior (*s*\*, *y*\*).

### 4.10 Numerical Examples

In this section, by numerical study, we illustrate the effect of different (linear, convex and concave) trade-offs on the level of the optimal backward help (*y*\*). We calculated the maximal long-term growth rate (fitness) of various populations as a function of *y* from the characteristic equation of the corresponding Leslie matrix. The value of *y* that gives the highest long-term growth rate termed as the optimal backward help (*y*\*). We also calculated the number of offspring and the offspring survival given the optimal *y*\*. We investigated the effects of different cost-benefit parameters on the evolvability of backward help (*y*). Life-history parameters are based on the figures from Mace (27). It is possible to generate all the possible combinations of cost-benefit trade-offs by setting the appropriate cost, benefit parameters to zero (*c, d, h*). Also, convex or concave cost-benefit functions can be achieved by setting the appropriate parameters (*c, d, h*) to smaller or to greater than one (see Table 2 for a summary of parameters). We used the following general Leslie matrix (see Figure 5 for a schematic description):

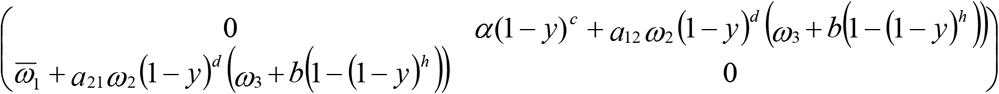

**Figure 5:**
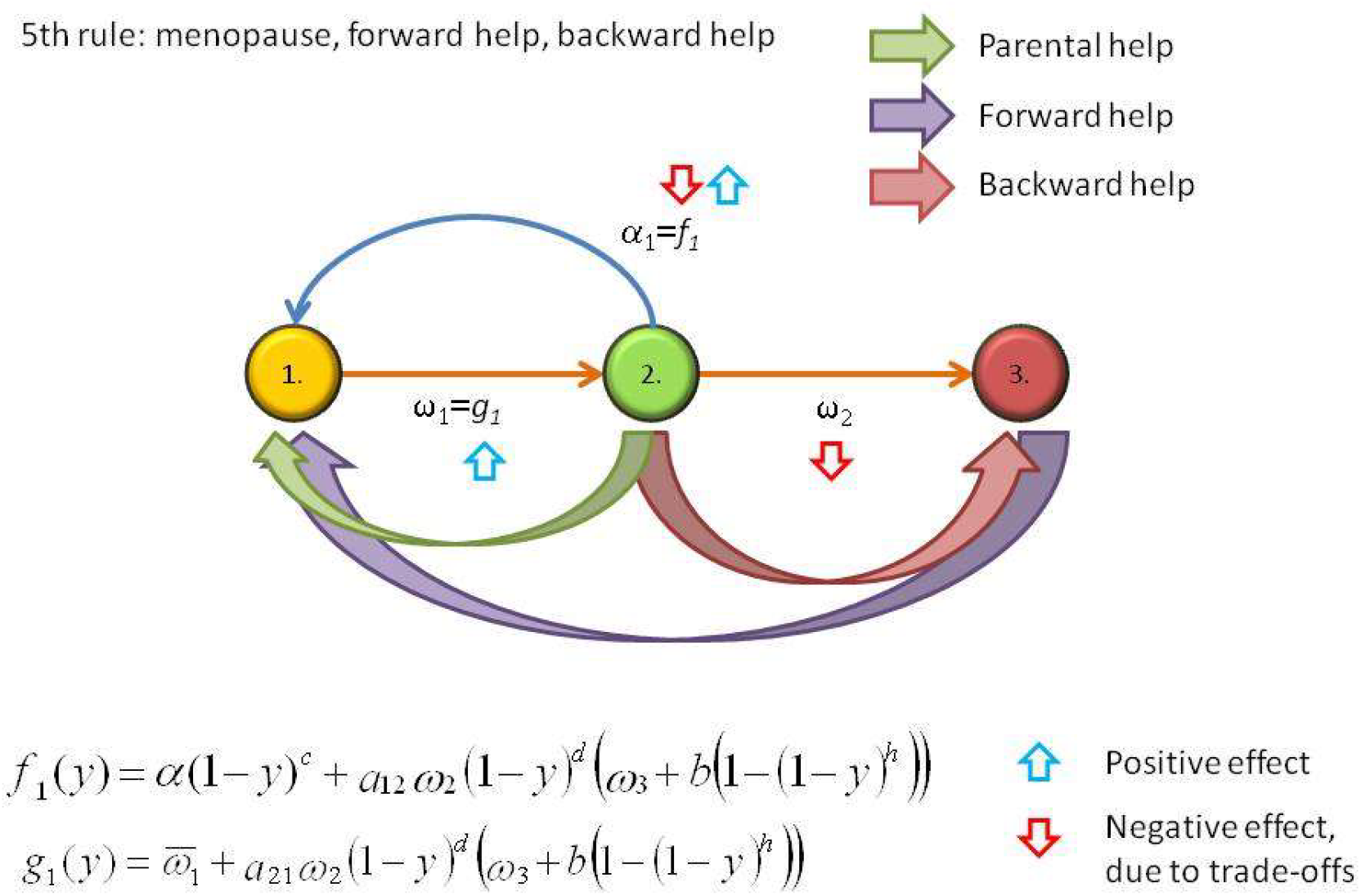
Schematic description of the two-age-class general model. Green arrows denote parental help; purple arrows denote forward help from the grandparents to the grandchild, finally red arrows denote backward transfer of resources from the parents to the grandparents.

**Table 2:**
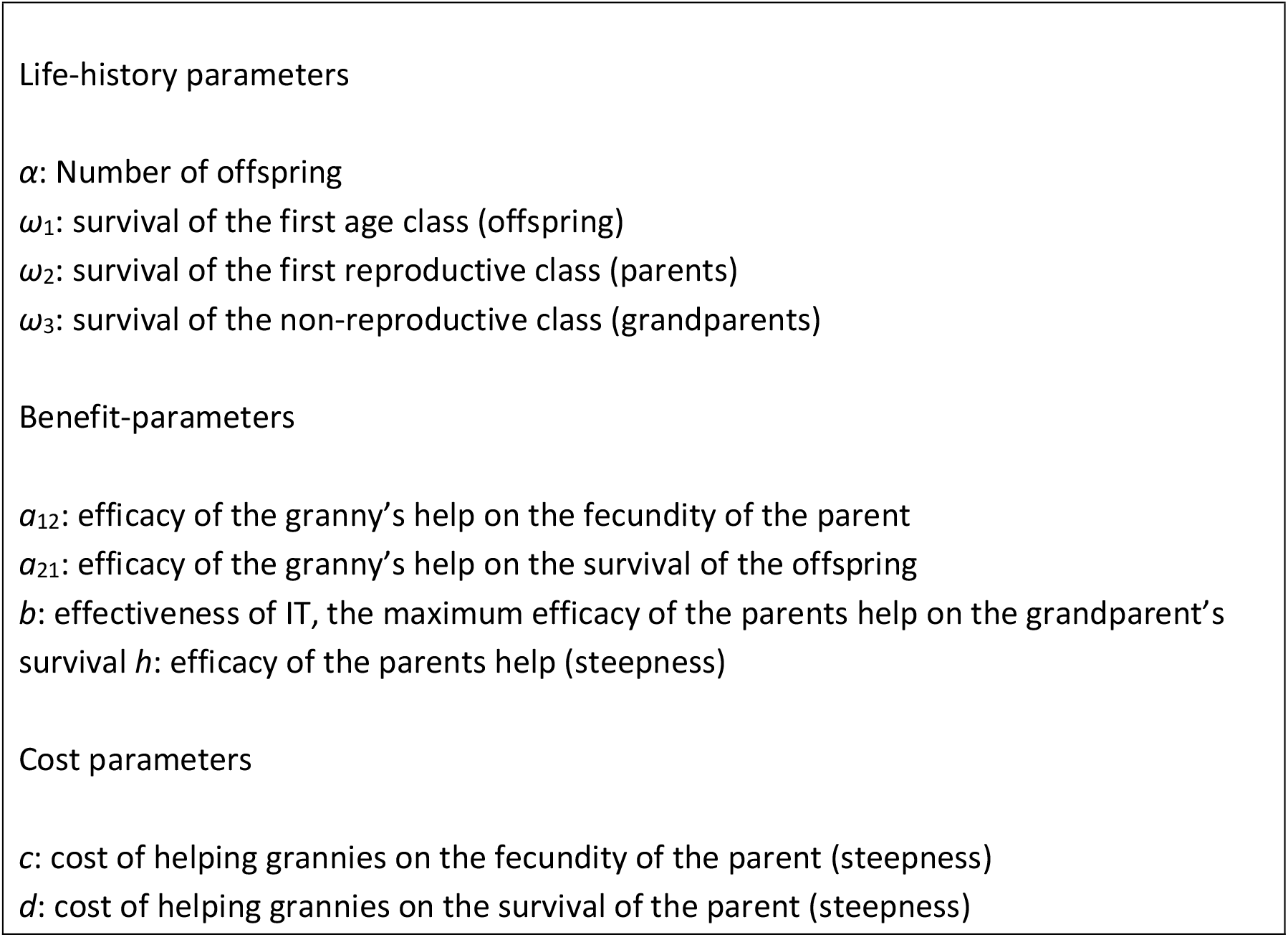
Parameters of the model.

Four possible combinations exist in terms of the benefit functions: (i) *a*_12_, *a*_21_ > 0; (ii) *a*_12_ > 0, *a*_21_ = 0, (iii) *a*_12_ = 0, *a*_21_ > 0; and (iv) *a*_12_, *a*_21_ = 0. In the first case, grandmothers give benefits for both the number of offspring and for the survival of them, in the second case they give benefit only for the number of offspring; in the third case they only give benefit for the survival of the offspring and finally, in the last case, they do not provide any benefit. This last case is not interesting for us, thus it will not be investigated any further.

In the same way, four possible combinations exist in terms of the cost functions: (i) *c, d* > 0; (ii) *c* > 0, *d* = 0, (iii) *c* = 0, *d* > 0; and (iv) *c, d* = 0. In the first case helping grandmothers imposes a cost on both the parents’ reproductive output and on the parents’ survival, in the second case only on the number of offspring, in the third only on the survival of the parent, and finally, in the last case it imposes no cost at all. Just as before, this last case is not interesting for us, thus it will not be investigated any further. See Table 3 for investigated parameter combinations.

**Table 3.** The investigated parameter combinations (see Figures S1-S7 for the corresponding results).

|  | Grandparental help | Shape of the cost function | Shape of the benefit function | Figure: |
| --- | --- | --- | --- | --- |
| 1. | $a_{12}, a_{21} > 0$ | $d = 1$ | $h = 1$ | S4 |
| 2. | | $d = 1$ | $h = 2$ | S5 |
| 3. | | $d = 0.5$ | $h = 2$ | S6 |
| 4. | $a_{12} = 0, a_{21} > 0$ | $d = 1$ | $h = 1$ | S7 |
| 5. | | $d = 0.5$ | $h = 2$ | S8 |
| 6. | $a_{12} > 0, a_{21} = 0$ | $d = 1$ | $h = 1$ | S9 |
| 7. | | $d = 0.5$ | $h = 2$ | S10 |

### 4.11 Illustrative numerical examples: results

IT evolves the most readily when the grandparental help increases both the survival of the offspring and the number of offspring (Figure S1-S3). Linear cost and benefit functions do not favour the evolution of IT (Figs. S1, S4, S6, *d*=1, *h*=1); conversely, convex benefit and concave cost functions promote the evolution of IT (Fig. S2, S3, S5, S7, *d*=0.5, *h*=2). It is possible to find cost parameters (*c, d*) where IT evolves even if the efficacy parental transfer and grandparental help (*a*_21_ and *b* respectively) is low (Figs. S2, S3). Conversely, it is possible to find (high) *a*_21_, *b* parameters where IT evolves even if it imposes a high cost on the survival of the parents or on the number of offspring (*d* and *c*, respectively, see Figs. S1, S2).

## Author contributions

J.G. and E.S. designed the study; J.G. and Z.V. analyzed the model; E.S., Z.V. and S.S. helped write the article, S.S. calculated the numerical examples and made the figures.

## Acknowledgements

We would like to thank István Scheuring and Tamás Czárán for helpful comments.

## Founding

J.G. and SZ.SZ. were supported by OTKA grant K 108974. E.S., J.G. and SZ.SZ. acknowledges support from GINOP 2.3.2-15-2016-00057 (Az evolúció fényben: elvek és megoldások); E.S. was supported by the European Research Council (ERC) under the European Community’s Seventh Framework Program (FP7/2007-2013) under ERC Grant Agreement 294332 (Project EvoEvo). SZ.SZ. was supported by the European Research Council (ERC) under the European Union’s Horizon 2020 research and innovation programme (grant agreement No 648693).

## Additional information

Supplementary Materials

